# Promoting diversity in ecological systems through toxin production

**DOI:** 10.1101/2023.03.06.531399

**Authors:** Ga Ching Lui, Sidhartha Goyal

## Abstract

Toxins are used by microbes to kill and compete, and hence, unlike cross-feeding from secondary metabolites, they are less often considered as a candidate to promote diversity in a microbial community. In contrast, some natural communities hint at the possibility that the presence of toxin genes and externally supplied antibiotics may support higher diversity. We address this gap with a model that shows how toxins in small and medium-sized communities can indeed promote diversity. The central idea of the model emerges from the combination of bacterial growth laws of resource partitioning and recent experiments suggesting a trade-off between toxin sensitivity and nutrient efficiency. The second puzzle in natural data is the presence of both negative and positive regulation of toxin production under similar environmental stress. Our model shows how this diversity in regulation of toxin production can promote toxin-producer fitness. In general, we provide a framework to integrate the role of toxins in microbial communities.

**Significance statement:** Structured organization of microbial communities into functional groups (consumers and producers) has been a framework to understand diverse populations. This framework leaves out the impact of toxin producers, which kill rather than provide the others with benefits, on diversification. Surprisingly, natural microbial communities show a correlation between the number of toxin genes in a community and its diversity. Our work shows how physiological constraints and trade-offs for individual bacteria can lead to a diverse, structured population where toxin-producers serve as a keystone species. This provides a pathway towards toxin production being established as a viable strategy in a diverse population: releasing toxins allows the survival of the less competitive strains, which would otherwise be driven towards extinction by competitive exclusion.

## Introduction

Cross-feeding on secondary metabolites released in a microbial community can change the composition of the population. Since producers of these metabolites promote the fitness of other species, cross-feeding can mediate commensal interactions between different bacterial species and sustain diversity under nutrient-limited conditions [17, 34, 37]. This further points to a possible coarse-grained description of community compositions by functional groups [15]. The core idea of this line of work is to consider these secondary metabolites as additional degrees of freedom that describe the system, allowing greater diversity at steady state. In other words, high diversity requires the production of a diverse spectrum of secondary metabolites.

Another category of secondary metabolites is toxins. Unlike cross-feeding, toxin increases the competitive advantage of toxin producers through allelopathy, meaning that toxins in the environment negatively affect their competing species [29]. This advantage can be mediated by interfering with the cellular processes of the competing species, such as suppressing protein biosynthesis, modifying and damaging cellular DNA, or by increasing the permeability of the cytoplasmic membrane, which effectively kills these cells [29, 45, 46, 18, 55]. In this sense, the production of toxins has an opposite effect on diversity compared to cross-feeding.

In models that describe interactions of species through toxins [22, 35, 19], it is common to observe multi-stability that leads to the dominance of a single species and lowers the diversity, which has been documented in early experiments such as [5]. In contrast, analysis of the Earth Microbiome Project dataset reports a correlation between diversity and antimicrobial toxin genes in rare taxa [33]. Furthermore, experimental work involving *Straphylococcus* spp. suggests that phenotypic diversity is promoted by intermediate antibiotic levels, and decreases at both high and low levels of antibiotics such as ciprofloxacin, amikacin, and streptomycin [28]. A related work on mouse guts shows that the presence of bile salt, which solubilizes lipids and is bactericidal, can drive coexistence in small systems with few mutants of *Escherichia coli* due to trade-offs between bile resistance and growth rate [9]. On the theoretical side, the work in [8] suggests that an autoinhibitor-producing species can stabilize a two-species system, and allelopathy can be established through two different paths: either relying on high initial abundances of the producers due to bi-stability, or the producer has to actively change the environment to reduce its own fitness and coexist with its competitor. The latter is described as “problematic from an evolutionary perspective” (or evolutionarily unstable), and one of the ways proposed to resolve this is to consider detoxification by the toxin-resistant species instead of toxin production [30]. It has also been suggested that toxin-producing phytoplankton may be responsible for maintaining coexistence through the effective coupling between species [48, 4].

Unlike cross-feeding, where secretion of secondary metabolites is not considered a cost to the secreting bacterium, the production of toxins is costly in terms of bacterial growth. This suggests that toxins should only be produced when necessary, and tight regulations may be in place to balance the benefit-to-cost ratio. One of the regulation mechanisms is quorum sensing [1], where toxin production is dependent on species abundance, and it has been shown mathematically to exhibit both pure states (only one species survives) and persistence (multiple species survive in oscillatory or chaotic attractors) [3]. Another potential mechanism is competition sensing [7], where toxin producers respond to abiotic stressors [39] and use environmental cues to infer ecological competitions. For abiotic stressors, there has been evidence that unfavorable growth conditions, such as low nutrient availability in the environment, contribute to an increased level of toxin production, and a rich body of literature has been dedicated to studying the effects of toxin production under different stressors [20, 27, 13, 7]. These effects guide the ecological dynamics and dictate the community structures. The intuition is that while species compete for these growth-limiting nutrients, toxins can increase the competitive advantage of the producer, but at a cost. Modeling suggests that species with constitutive production of toxins tend to be out-competed by species with regulated production through quorum sensing or competitive sensing on abiotic stressors [39].

In this work, we present theoretical work that shows how toxins can counterintuitively play a role in supporting diversity. Suppose that the environment of a particular system can be described by some low-dimensional space such that for each species, the space can be partitioned into two different regions: (1) bacterial growth rate greater than removal rate, and (2) negative net growth. The two regions are separated by a curve plotted in Figure 1A, which is called the zero net growth isocline (ZNGI). ZNGI denotes a set of environmental conditions that correspond to the steady states when that species is present in the system:

**Figure 1.**
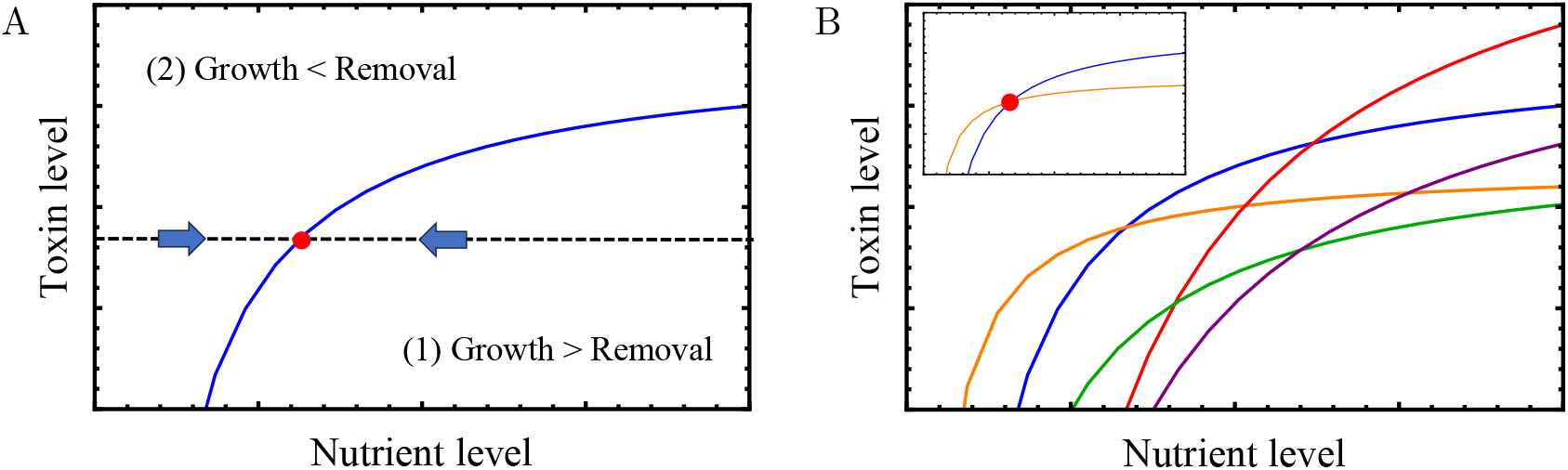
(A) Environment defined by nutrient and toxin levels with a single species. In region (1), where the toxin level is low or the nutrient level is high, the growth rate of the species is larger than the removal rate, leading to large consumption of nutrients. In region (2), hostile environments lead to a small population size, and the overall nutrient level is replenished by the external supply. Due to this feedback, the nutrient and toxin levels at steady state are given by the ZNGI in Eq. (1). (B). The inset shows the ZNGIs of two species, which can coexist at the nutrient and toxin levels given by the red dot. For multiple species, it is difficult for all ZNGIs to intersect, and coexistence is not expected.

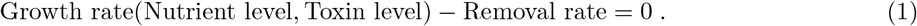

If another species is added to the system, they can coexist only at the intersection of the ZNGIs (inset of Figure 1B). Now, if more species are added to the system as shown in Figure 1B, either through mutation or immigration, it is improbable that all of these ZNGIs intersect in this low-dimensional space. In this paper, we propose two different paths through which high diversity can be facilitated by toxins: (1) imposing symmetry to the entire population through trade-offs and (2) adjusting the ZNGIs by changing the rate of toxin production. We then address under what circumstances is the strategy of toxin production evolutionarily stable. Finally, we introduce regulation by considering the sensing of abiotic factors (nutrient or toxin levels) and show that persistence through regulation can also increase diversity.

## 1 The model

### A. Growth rate functionals respecting proteome constraint

To investigate if toxins can play a role in sustaining diversity by trade-offs and by the increase in the number of abiotic factors, we need an approach that organically incorporates nutrient competition, toxin susceptibility, and metabolic constraints into the modeling of bacterial growth. We adopt recent models [57, 50] that describe metabolic constraints in terms of proteome partitioning, where the bacterial growth rate 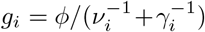 for species *i* depends on nutrient efficiency *ν*_*i*_, translation efficiency *γ*_*i*_ in synthesizing proteins, and is subjected to the same proteome fraction devoted to growth (denoted by *ϕ*) for all species in the system. In SM Sect. 1.A., we show that in a system with one growth-limiting nutrient and one toxin, the bacterial growth rate for species *i* can be written as

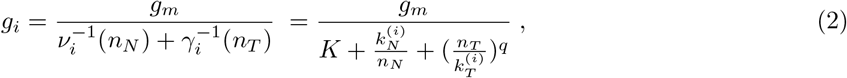

where 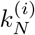 and 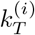 are the Michaelis-Menten constants that determine the scales of nutrient level *n_*N*_* and toxin level n_*T*_ in the environment, and *q* > 0 is the Hill coefficient for toxin susceptibility. The parameter *K* is a constant independent of nutrient and toxin levels, such that *g_*m*_/K* is the maximal growth rate in an optimal environment with an abundance of nutrients at a low toxin level when 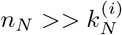 and 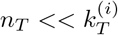, and is proportional to *ϕ*.

Suppose that a certain fraction *f*_*i*_ <1 of the nutrient flux is redirected from bacterial growth towards toxin production (see SM Sect. 1.B), there is an associated cost incurred in the nutrient efficiency *ν*_*i*_, such that

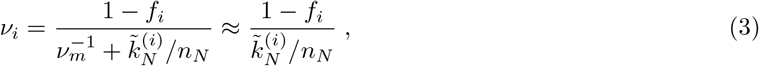

in the limit of a nutrient-depleted environment 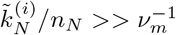. Here, ν_*m*_ is the maximum efficiency when the nutrient is abundant, *f*_*i*_ is the production fraction and 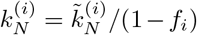 depends on cost of production: 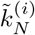 is the Michaelis-Menten constant when no toxin is produced. By substitution of Eq. (3), the growth rate in Eq. (2) becomes

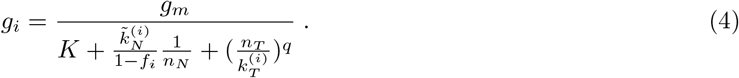

As the toxin production fraction *f*_*i*_ increases, the bacterial growth rate *g*_*i*_ decreases due to a decrease in the nutrient efficiency *ν*_*i*_.

### B. Competition in a well-mixed continuous culture system

Populations in continuous culture systems are expected to be reasonably well-mixed with no appreciable spatial structures [56]. In laboratories, continuous long-term culturing is achievable with automated real-time monitoring of population compositions at an affordable cost [41]. As such, we focus on ecological diversity in a chemostat system, which is a continuous culturing device that pumps in fresh medium and removes stale medium at some constant rate. Chemostat experiments allow easy control over population sizes and environmental conditions, such as nutrient levels in the culture, with less fluctuation introduced externally. This provides a reasonable model for natural populations from gut microbiota to sewage [9, 31], although these have spatial structures that may also play a significant role [10, 54]. In general, a chemostat system is described by the following set of equations:

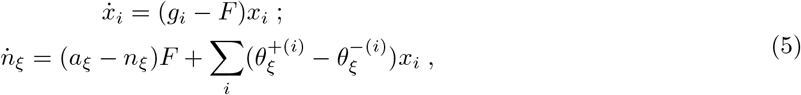

where *x*_*i*_ is the abundance of species *i*, *g*_*i*_ is the bacterial growth rate and *n*_*ξ*_ is the concentration of some abiotic factor *ξ*, with *ξ* = *T* for the toxin and *ξ* = *N* for the nutrient. A fresh medium, with nutrient and toxin concentrations *a*_*ξ*_, is pumped into the chemostat at a flow rate of *F*. The production and consumption of abiotic factor *ξ* are denoted by 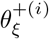 and 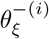, which describe the change in nutrient and toxin levels due to species *i*. By adjusting the flow rate *F* of the chemostat and the supply concentrations *a*_*ξ*_ of the abiotic factors, we can alter the selective pressure imposed on the species and modify the composition of the community.

For the remainder of the paper, we consider various systems defined by Eq. (5) with different forms of growth rates *g*_*i*_ and rates associated with abiotic factors 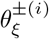. To survive in the chemostat at steady state, bacteria must change the levels of abiotic factors so that the bacterial growth rate is equal to the removal rate due to the flow of the medium in the chemostat, or *g*_*i*_(*n*_*N*_, *n*_*T*_) = *F*. We first consider the constitutive production of toxins, before investigating the effect of regulation on diversity. Our system is summarized in Figure 2.

## 2 Results

### A. Equalization by trade-offs and stabilization by toxin production

Can toxins, which are primarily produced to kill, increase diversity at steady states? We consider the simplest scenario of having only one growth-limiting nutrient *N* and one toxin *T* in the system, with concentrations *n*_*N*_ and *n*_*T*_. The system is defined by Eq. (5) with bacterial growth rate given in Eq. (4). Nutrient *N* is externally supplied through the inflow of the fresh medium with concentration *a*_*N*_ > 0, and is consumed by species i at a rate of 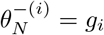 per cell to support either bacterial growth or toxin production. Toxin *T* may or may not be externally supplied with *a*_*T*_ *≥* 0 and is produced by the species at a rate of 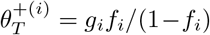, which increases with production fraction *f*_*i*_. In our model, the toxin is not consumed and the nutrient is not produced by the cells 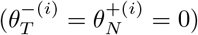. As *f*_*i*_ increases, the rate of toxin production 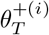 increases at the expense of a decreasing growth rate described in Eq. (4). We show the derivation in SM Sect. 1B.

Suppose that in this system there are five species (*i* = 1, …, 5), each with the Hill coefficient *q* = 1 for toxin susceptibility in Eq. (4), what are the potential outcomes of the competition? According to the competitive exclusion principle [21], when there is only one externally supplied growth-limiting nutrient (with toxin supply concentration set to be *a*_*T*_ = 0, and that there is no cross-feeding), it is not possible to have more than one species survive at steady states. The fittest species would dominate the entire population and drive other species towards extinction. However, we notice that if there is at least one species that produces toxins, and that the following condition holds:

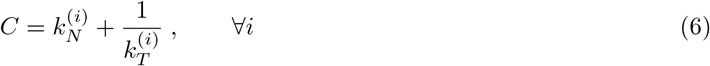

where *C* > 0 is some constant, then the system can be brought to a state where all five species coexist. We will show in the next subsection and in SM Sect. 4 that Eq. (6) can be understood as a trade-off between the nutrient-driven growth rate and the toxin sensitivity observed in experiments. For the time being, we focus on the consequences of Eq. (6) and consider species *i* = 1 to be the only toxin producer in the system.

Given this trade-off, the amount of toxins released by the producer determines the competition outcome, and diversity is sustained only at intermediate values of production fraction *f*_1_. Figure 3A shows the species abundances at steady state when *f*_1_ changes, and illustrates three different regimes. At small *f*_1_ < *f*_*c*1_ in zone (1), the system shows a pure state where the toxin producer dominates the population. In zone (2) with *f*_*c*1_ < *f*_1_ < *f*_*c*2_, the producer only accounts for a portion of the total biomass, while other species successfully invade the population. Diversity increases while the total biomass sustained in the system decreases compared to that in zone (1). At large *f*_1_ > *f*_*c*2_ in zone (3), the system returns to a pure state that shows bistability: either the toxin producer or one of the non-producers survives.

**Figure 2.**
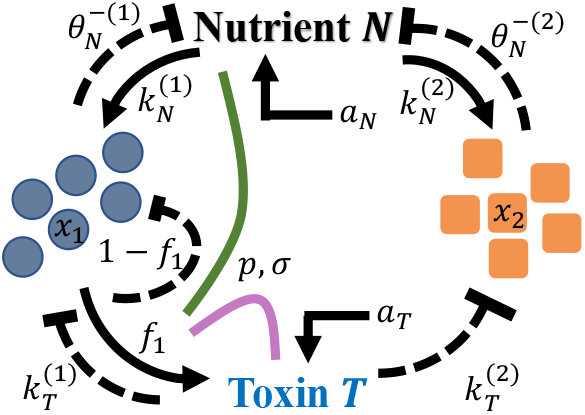
Diagram showing a system with a toxin non-producer (orange) and a toxin producer (blue), described by Eq. (4) and Eq. (5). Toxin production fraction *f*_1_ is constant under constitutive production in Sect. 3.1. Regulation in Sect. 3.2 changes *f*_1_ under nutrient sensing (green line) or toxin sensing (purple line), which are parametrized by *ρ* and *σ* in Eq. (9).

**Figure 3.**
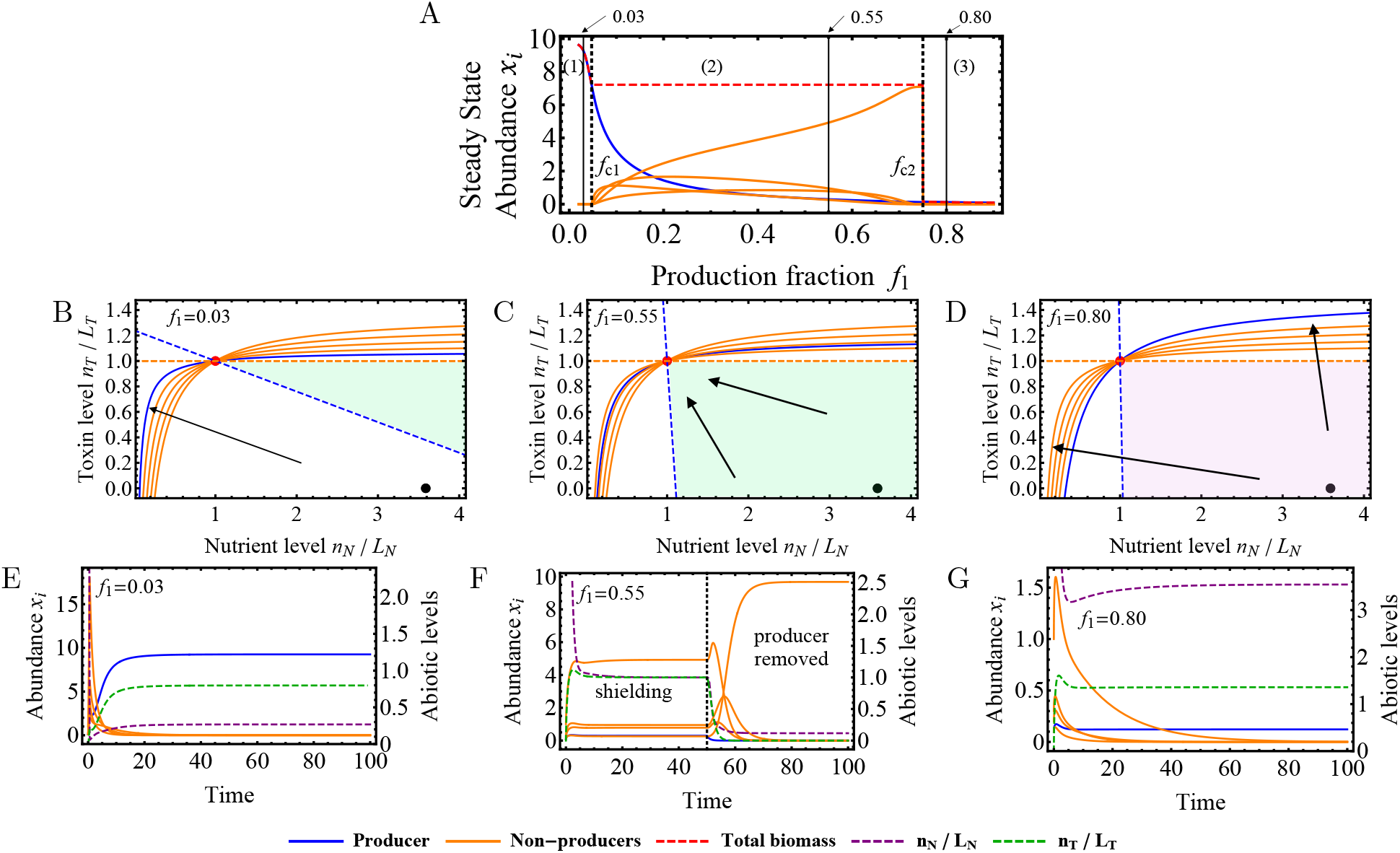
Diversity sustained by constitutive toxin production. (A) Species abundances and total biomass as production fraction *f*_1_ changes. The three values *f*_1_ = 0.03, 0.55, 0.80 are marked by solid black lines showing three phases, and the two transitions are at *f*_1_ = *f*_*c*1_, *f*_*c*2_, separating the range of *f*_1_ into zone (1), (2) and (3). (B-D) ZNGIs (solid) and impact lines (dashed) in abiotic space for the producer (blue) and the non-producers (orange). The nutrient level *n*_*N*_ and toxin level *n*_*T*_ are scaled by constants 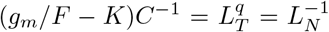 (SM Sect. 2). If the supply vector (*a*_*T*_ /*L*_*T*_,*a*_*N*_ /*L*_*N*_), notated by black dot, lies in the shaded region, the coexistence solution is physical, which can either be stable (green) or unstable (magenta). Black arrow shows directions of the dynamics, affecting the stability of coexistence. (E-G) Time series of species abundance (solid) and abiotic factors levels (dashed). In (F), the producer is removed at *t* = 50, leading to the collapse of the community.

A necessary condition for coexistence is fitness equalization. Consider the ZNGI to be *g*_*i*_(*n*_*N*_, *n*_*T*_) − F = 0. We plotted the ZNGIs of the five species in Figure 3B-D for the three different zones. Notice that due to the symmetry imposed by the trade-off in Eq. (6), all ZNGIs intersect at 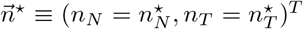: this denotes the environmental condition in which all species are equally fit and have a growth rate g_*i*_ equal to the flow rate *F*. This is in contrast to the case without trade-offs in Figure 1B, where 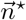 does not exist. When there is no external supply of toxins (*a*_*T*_ = 0), the presence of a toxin producer is required to bring the toxin level up to 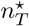 to achieve fitness equalization.

Under what circumstances then is the toxin producer able to create an environment in which all species are equally fit? To describe how species *i* changes the environment through both the consumption and production of toxin *T* and nutrient *N*, we define the impact factors 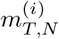 as

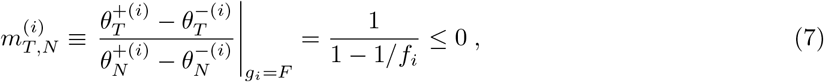

which turns out to be uniquely determined by the production fraction *f*_*i*_. We further define the supply vector 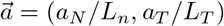 and the “impact lines” 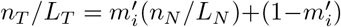 with slope 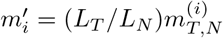, where *L*_*N*_ and *L*_*T*_ are constants that provide proper scalings, and plotted them in Figure 3B-D. We show analytically in SM Sect. 2 that coexistence can only be sustained when the supply vector 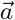 lies within the shaded region bounded by the impact lines. For toxin producers, the impact lines always have a negative slope 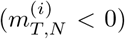 as nutrient *N* is always consumed and toxin *T* is always produced per unit cell growth; a species that does not produce toxins would have a horizontal impact line (or 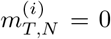). Without any toxin producer, the bounded region has zero area and coexistence is not possible unless the supply vector 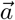 is fine-tuned: the supply concentration of toxin *a*_*T*_ has to match exactly with 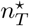. The presence of a toxin producer provides an upward force to the flow in the abiotic factor space, so that fine-tuning is no longer required. As *f*_1_ increases, the impact ratio 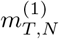 of the producer increases, which widens the bounded area. Therefore, coexistence can be sustained at a lower nutrient supply concentration *a*_*N*_. This gives the first transition at *f*_1_ = *f*_*c*1_ that we observe in Figure 3A, where 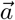 is on the impact line of the toxin producer, or 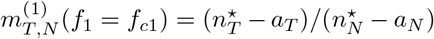. Figure 3E-F shows the time-series of both species abundances and abiotic levels before and after the transition. After the transition, we notice that an increase in diversity is accompanied by a matching of toxin level with nutrient level *n*_*T*_ /*L*_*T*_ = *n*_*N*_ /*L*_*N*_ = 1. We call this the shielding effect: the microbiome self-organizes to shield off the asymmetry in the supply concentrations *a*_*N*_ ≠ *n*_*T*_, such that the rescaled nutrient and toxin levels can be treated as equals. Therefore, strategies preferential towards nutrient competition are equally optimal to strategies that favor toxin resistance. This was previously observed in models of competition over alternate resources [52, 43, 32] in large randomized systems [53], but not in toxin models. Shielding effect is sustained only in the presence of the producer; removing the producer from the population leads to the collapse of diversity in the population, as shown in Figure 3F.

The implication is that the producer affects the stability of the coexistence state. As the production fraction *f*_1_ increases, both 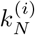 and 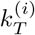 decrease due to Eq. (6) and the ZNGI is shifted. The ZNGI for the producer at large 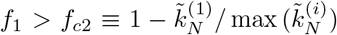 is shown in Figure 3D. Even though the supply vector 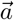 remains in the bounded region, the coexistence state is no longer stable, as shown in SM Sect. 3. The time-series in Figure 3G shows the breakdown of the shielding effect as toxin level and nutrient level separate, resulting in the sole survival of a single species. As *f*_1_ approaches 1, the toxin-producing species can no longer be sustained as its growth rate decreases to zero, as the nutrients consumed are funneled into the production of toxins.

### Trade-offs

Do we see the above hypothesized trade-off between nutrient efficiency and toxin susceptibility in natural systems? Just to reiterate, the trade-off required for fitness equalization suggests that strains that are more efficient at consuming nutrients pay a price by being more susceptible to toxins. We went further and postulated a specific functional form for this trade-off. As potential evidence for this trade-off we first present the experimental data from [42] that utilized multiple *E. coli* strains in Figure 4A. The data plotted shows the growth rate of a strain relative to that of a reference strain in the absence of toxins against a strain’s resistance to three different antibiotics. We show in SM Sect. 4 that the relation

**Figure 4.**
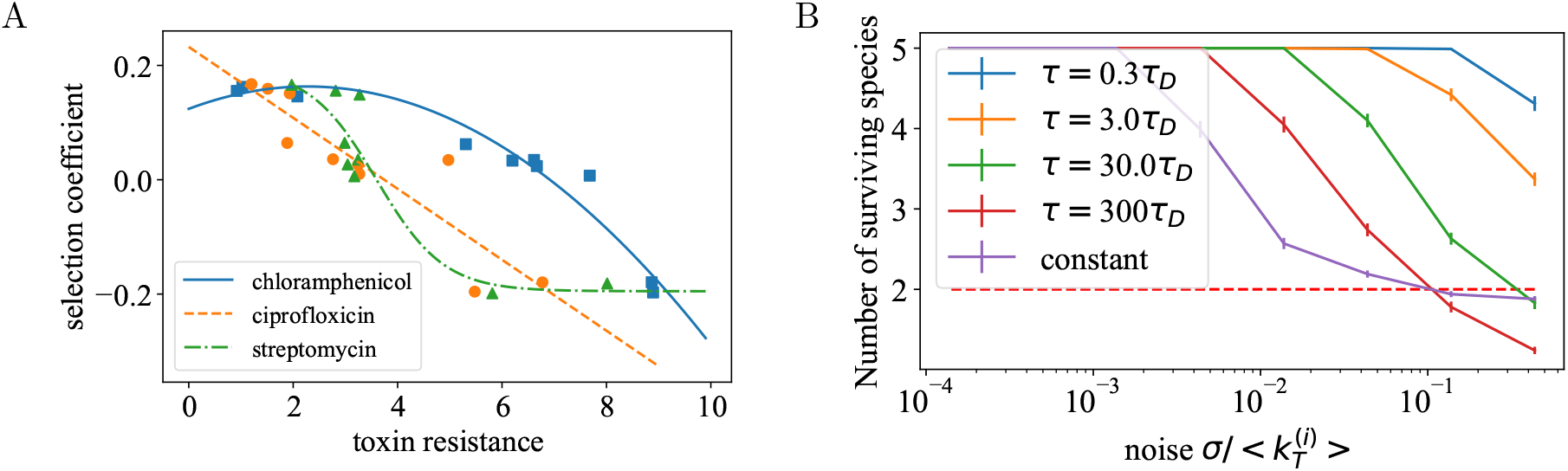
(a) Data adapted from Ferenci et al. [42] to illustrate trade-offs, showing three different trends of nutrient competence (measured by selection coefficient) as toxin resistance increases. These trends (accelerating, linear and decelerating) are qualitatively captured by the model. (b) The number of surviving species as *σ* and switching time *τ* changes. Red, dotted line notates the upper limit of diversity given by the competitive exclusion principle. Parameters are given in SM Sect. 8.

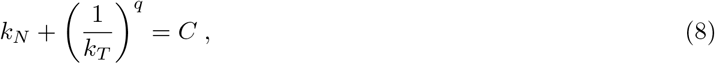

with *k*_*N*_, *k*_*T*_, C being the fitting constants, can recapture the different trends between the relative growth rate and toxin resistance shown in Figure 4A. Generally, the shapes of trade-offs affect the fitness of a species, and a convex (or concave) trade-off makes it more beneficial to be a specialist (or a generalist) [12]. In contrast, when the environmental condition (*n*_*T*_ /*L*_*T*_)^*q*^ = *n*_*N*_ /*L*_*n*_ = 1 is satisfied, the shielding effect can be achieved in our model, such that all species (including both specialists and generalists) attain the same fitness, regardless of the values of *q* or the shapes of the trade-offs. It is clear that Eq. (6) in Sect. 2.A is a special case of Eq. (8) corresponding to *q* = 1. While Figure 4A shows that data generally follow the trend due to the trade-off, it also shows that data can depart from the exact relationship, and therefore from Eq. (8). This brings a natural question: how fine-tuned does the trade-off have to be for fitness equalization?

To answer this question, we considered a system with five species characterized by the Michaelis-Menten constants 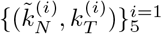 which satisfy the trade-off in Eq. (6), but here we made these parameters noisy in time: 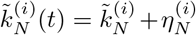 and 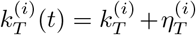. The deviation from the trade-offs may happen on long time scales due to mutations that lead to changes in the phenotype, or on short time scales due to fluctuations in gene expression. To test the time dependence, we sampled the Michaelis-Menten constants 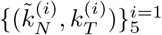 across different timescales *τ*. The noise terms 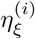 were sampled from normal distributions *𝒩*(0, *σ*^2^) for every *τ* units of time. The number of surviving species after transient dynamics is recorded over an ensemble.

For fast switching time, i.e. *τ* ≪ *τ*_*D*_, where *τ*_*D*_ is the cell division time, the diversity is most robust to noise; for example, even for *τ* = 0.3*τ*_*D*_ the diversity remains unchanged until the noise gets beyond ~20%. The observed diversity falls as the switching time *τ* increases. By *τ* ~ 300*τ*_*D*_, a 10% noise is enough to destroy the shielding effect completely. Overall, this suggests that while mutations can break the equalization scheme, fitness equalization is quite robust to gene expression fluctuation.

### B. Toxin regulation leads to new behavior - persistence, supersaturation, and producer’s improved survival

We showed above how trade-offs sustain diversity through equalization under constitutive production. However, toxin production is costly for bacterial growth, and a more effective strategy is to avoid producing toxins unless necessary. Indeed, up-regulation of toxin synthesis in response to various stressors has been reported [7]. One way to model this is to assume that a species produces more toxins when the growth rate is slow [35], which is indicative of a stressed environment. Here, we explicitly model the dependence of toxin regulation on environmental cues by treating the toxin production fraction as a function of either the nutrient level *f*_1_ = *f*_1_(*n*_*N*_) or the toxin level *f*_1_ = *f*_1_(*n*_*T*_), as shown in Figure 2. Consequently, having a generic trade-off in the form of Eq. (8) is no longer possible. Without regulations, the expectation is that no more than two species can coexist in a system with only two growth-limiting factors [36], namely the externally supplied nutrient and the produced toxin.

To understand this better, we considered the competitions between a single toxin producer (*i* = 1) with one (*i* = 2) or more (*i* ≥ 2) toxin non-producers, with abundances *x*_*i*_. Specifically, the bacterial growth rates in Eq. (4) and the dynamics for both the nutrient and the toxin levels in Eq. (5) for this system can be explicitly rewritten as

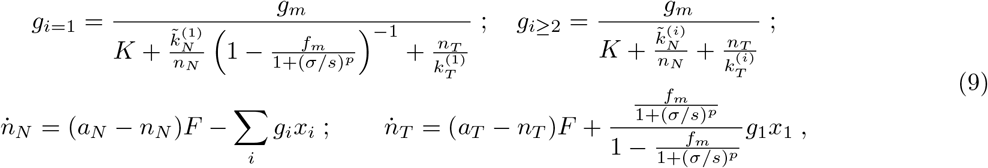

where *K* is a constant unrelated to nutrient and toxin levels, such that *g*_*m*_/*K* is the maximal growth rate in an optimal environment rich in nutrients and low on toxins. We model the dependence of the toxin production fraction from Eq. (4) as a Hill function, *f*_1_ = *f*_*m*_/[1 + (*σ*/*s*)^*p*^], where s = *n*_*N*_ for nutrient sensing and *s* = *n*_*T*_ for toxin sensing. The Hill coefficient *p* determines how acute the response is, and can take both positive and negative values to span both positive and negative regulations. The constant *σ* sets the concentration scale of s for sensing. And finally, the parameter *f*_*m*_ is the maximum value that the toxin production fraction *f*_1_ can take; note that *f*_*m*_/2 is the value of the production fraction without regulation.

In this section, we start with a two-species system in which the parameters are set such that the trade-off is intact under the constitutive production of toxins, but now toxin production is regulated. We saw that regulation changes the behavior of the system through altering the fitness of the toxin producer. As shielding is no longer in effect, only certain choices of *p* and *σ* would allow the toxin producer to thrive. To further understand the role of regulation for a community, we then considered cases where coexistence is infeasible without regulation; either because the species do not share any trade-offs or the state of fitness equalization is unstable. Lastly, we explored how regulation may enhance the diversity beyond what is supported by the environment - called supersaturation.

### Regulation breaks shielding effect and can hurt the producer’s ability to survive

How does regulation change the system that exhibits high diversity sustained by trade-offs under constitutive toxin production? Following Eq. (6), we set the Michaelis-Menten constants to respect the trade-off: 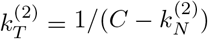 and 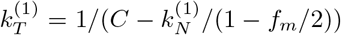. As the toxin production fraction *f*_*m*_ increases, the toxin producer is more resistant to the toxin. Since here we are only considering 2 species, trade-off is not needed for the two ZNGIs to intersect, as shown in the subset of Figure 1B. But we use it to relate the behaviours with no regulations, or *p* = 0, studied in Sect. 2.A. Depending on *f*_*m*_ then, the two species can show coexistence or bistability under constitutive production (*p* = 0), which correspond to zones (2) and (3) in Figure 3 respectively. We did not consider zone (1) as the change in regulation does not lead to transitions: the producer is always the fittest, with or without regulation.

We focus here on discussing the results for nutrient sensing, for it will become evident later in this section and in SM Sect. 6 that similar behaviours are also exhibited by toxin sensing. Figure 5A-B show the phase diagrams as we span the Hill coefficient p and scaling constant σ. For small |*p*| the behavior is as expected, but as |*p*| increases, the two species no longer share the trade-off between the effective nutrient efficiency 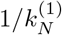 and the toxin resistance 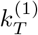. This broken symmetry in competitiveness of the two species and can lead to pure states dominated by either the toxin producer (T) or the non-producer (NP), which signifies a change in fitness due to sensing. Note that in the cases shown in Figure 5A-B, there is some significant phase space where the toxin producer loses out due to regulation.

**Figure 5.**
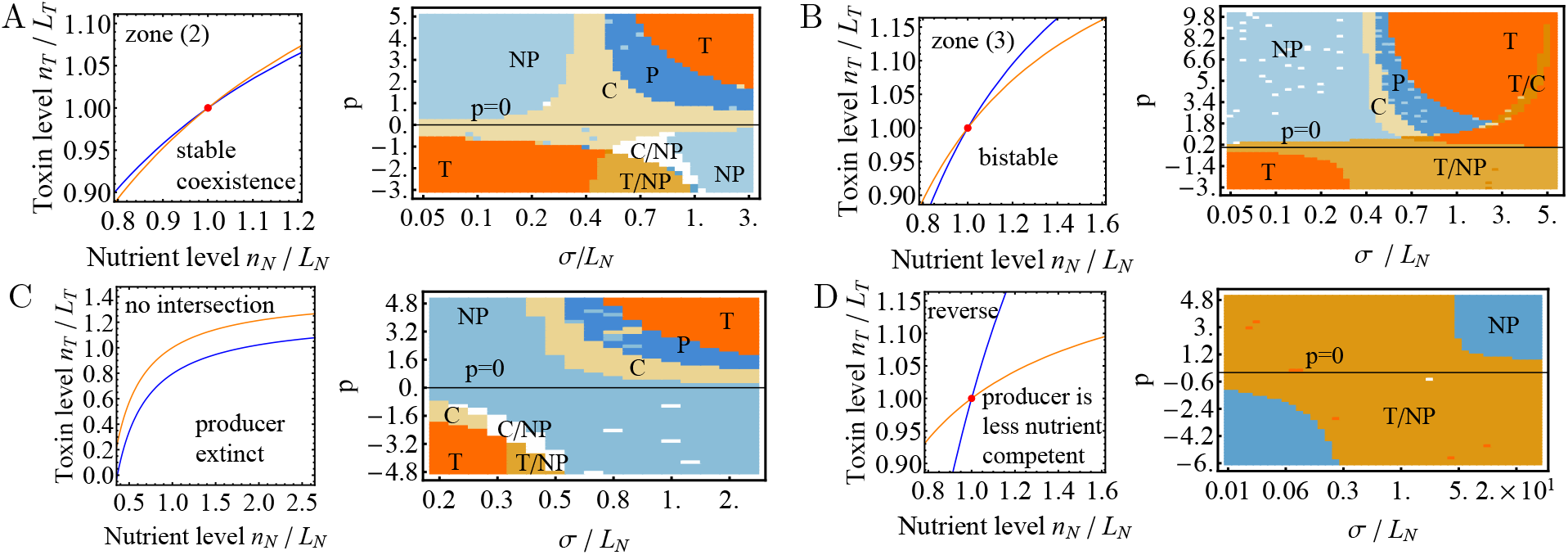
Regulation through nutrient sensing. The four regimes under consideration are: (A) small *f*_1_ = 0.55 in zone (2); (B) large *f*_1_ = 0.9 in zone (3); (C) no intersections of ZNGIs; and (D) the reverse case for nutrient efficiency. All except (D) has 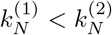, with ZNGIs similar to Figure 3D. Left figure shows ZNGIs of the producer (blue) and non-producer (orange) at p = 0. Right figure shows the phase diagram as Hill function p and sensing scale *σ* vary. *L*_*N*_ is a constant that sets the scales. Possible competition outcomes are coexistence (C), persistence (P) and pure states dominated by producer (T) or non-producing species (NP). Slash notates bistability of two outcomes (T/C,C/NP,T/NP). Parameters are given in SM Sect. 8.

### Regulation rescues species with large toxin production fraction by increasing diversity

If toxin regulation can be detrimental to a toxin producer, as we have argued above, why do they choose to regulate the toxins then? To understand this better, we considered the situations where constitutive production was indeed too expensive and not beneficial to the producer. Here, we found that regulation provides a path for the producer to establish itself in the community.

For Figure 5C, we changed the Michaelis-Menten constants such that the toxin producer without sensing (*p* = 0) is always driven to extinction by the non-producer in the blue regions (NP) due to the cost of toxin production. Note that in the absence of the trade-off, the ZNGIs of the two species do not intersect under constitutive production: the non-producer can tolerate a higher toxin level and a lower nutrient level than the producer without regulation, and is therefore the fitter species. Here, nutrient sensing allows producers to survive in the regions of the phase diagrams that correspond to pure state (T), coexistence (C) or persistence (P). These regions in which the producer becomes more competitive either correspond to large *σ* with *p* > 0 or small *σ* with *p* < 0. We show in SM Sect. 5 that both positive and negative sensing shift the ZNGI of the producer by increasing its fitness, leading to coexistence of the two species.

In the presence of the trade-off, a producer with a large production fraction *f*_*m*_ will be able to invade under constitutive production only if its initial abundance is high. As shown in Figure 5B, regulation can move the system away from the brown region (T/NP), allowing the producer to gain fitness and remain within the population regardless of initial conditions.

### Toxin down-regulation promotes toxin-producer fitness

When the toxin producer is more nutrient-efficient than the non-producer 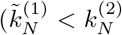, ignoring the cost of toxin production), we observed a common driving force that acts on the competitiveness of the producer for both positive sensing (*p* > 0) and negative sensing (*p* < 0), as seen in regions where toxin producers are the sole survivors in top-right and bottom-left of Figure 5A-C. With positive sensing, the top-right corner corresponds to the limit where toxin production is down-regulated most of the time unless the toxin level is high. Interestingly, we also see a similar trend with negative sensing(*p* < 0), where in the bottom-left corner the toxin production is again mostly down-regulated once the nutrient level exceeds some small threshold *σ*. Hence, for both sensing schemes: positive or negative, the producer thrives when it mostly down-regulates its toxin production over a wide range of values of nutrient levels n_*N*_ (*p* > 0, very large *σ* and *p* < 0, very small *σ*).

It might make sense for a species to dedicate some of the incoming nutrients to toxin production if it is one of the nutrient-competent species. But what if the producer is less nutrient-competent than the non-producer: 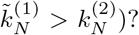 Based on the observed trade-off, the producer would be more toxin resistant than the non-producer 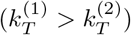. This is the regime in which previous studies show that diversity is not sustainable under constitutive production [8]. We call this the reverse case, which is plotted in Figure 5D. At *p* = 0, either the producer or the non-producer, but not both, survives at the steady state. Regulation does not change the diversity of the system. Since the non-producer is more susceptible to the toxin, it is fitter when the toxin production is down-regulated, and dominates the population at either large *σ* for positive sensing and small *σ* for negative sensing.

### Regulation allows for persistence and supersaturation

Beyond the steady-state behaviours we have discussed above, regulation also allows for persistence where the abundances of the different species oscillate. Notably, this is observed only for positive regulation *p* > 0. With positive sensing, a high level of nutrient or toxin (which corresponds to a fast growth rate) up-regulates toxin production, and thereby the production cost for the producer. This, in turn, lowers the producer’s growth rate. Essentially, this provides negative feedback on the toxin producer and creates a possibility of oscillations. In contrast, during negative sensing (*p* < 0), a higher level of nutrient or toxin down-regulates toxin production and hence increases the growth rate, which serves as positive feedback. Not surprisingly, we see more bistable regions in this case.

Persistence then provides another pathway to increase diversity beyond what is supported by the environment, called supersaturation. It has been shown in models that supersaturation can be generated when the population competes for more than three nutrient resources [24], and that the system observes certain symmetries in the nutrient efficiencies [23]. To this end, we considered the system defined by Eq. (9) with one producer (*i* = 1) with positive sensing and multiple non-producers (*i* = 2, 3…). Without trade-offs, we should expect at most two coexisting species at steady states [36, 21], as discussed above. By comparison, Figure 6A shows that a producer growing along with two non-producers, even though the ZNGIs of the three species in Figure 6B share no intersections. However, this is not a steady state, and here regulation of toxin production leads to oscillating levels of the nutrient and the toxin, preventing any of the species from being consistently fit to dominate the population.

**Figure 6.**
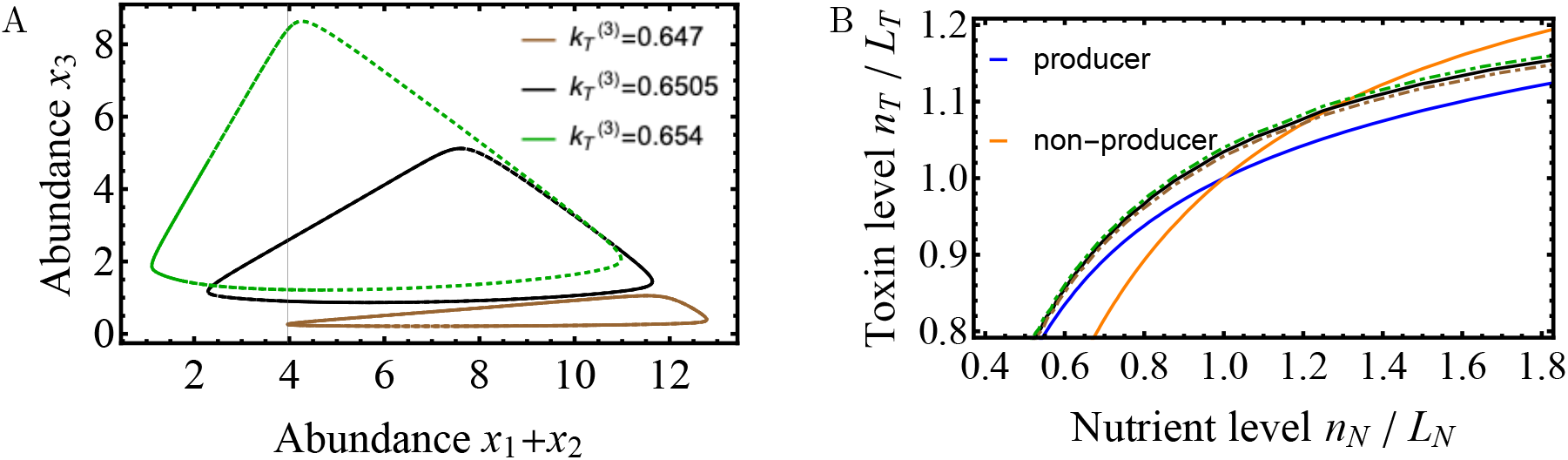
Supersaturation of three species supported by a toxin and a nutrient source due to negative feedback under nutrient sensing. A third species with varying 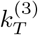 (Green, black and brown) is added to a two-species system (blue, orange). (A) Limit cycles of species abundances after transient dynamics. Supersaturation is not sustainable outside of this range. (B) ZNGIs of the three species. Slight changes in toxin resistance of the third species greatly affect the fitness and the competition outcome.

This pathway towards diversity, however, is limited in our model. First, small differences in the parameters drastically change how the system behaves and can break supersaturation. We kept all parameters constant while varying the toxin efficiency of the third species 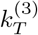 within a small range (~1%), with no discernible qualitative differences in the ZNGIs plot in Figure 6B, and plotted the species abundances in Figure 6A. As 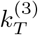 decreases within this range, which makes the species less competitive, the maximum abundance attained by the third species drops (from the green curve to the black and brown curves). Supersaturation is not sustained outside of this limited range. Whether there is a trade-off that may allow this “fine-tuning” of parameters for species to be achieved remains to be explored. Notice, however, that given the presence of the “fine-tuned” species, the setup of the system does not need to be fine-tuned, and supersaturation can be sustained over a range of flow rates and nutrient supply concentrations (see SM Sect. 7A).

Second, this mechanism is not scalable in our model. The parameter range that supports supersaturation quickly dropped from ~ 1% to ~ 0.1% when another species was added to the system (see SM Sect. 7B), and we have not been able to pack more than four species. If supersaturation is to be a significant driving force for diversity in nature, there is likely to be additional symmetries (e.g., trade-offs) at play. While oscillation is frequently observed in systems involving toxins, for example, in ecological suicide [6, 44] where a toxin producer releases autoinhibitors into the environment that lead to oscillating population sizes, its roles in supersaturation remain elusive.

Finally, qualitative behaviours of how regulation under nutrient sensing (*s* = *n*_*N*_) are also observed in toxin sensing (*s* = *n*_*T*_). In SM Sect. 6, we show that both sensing schemes share a common mechanism: they effectively regulate toxin production based on the growth rate of the toxin producer 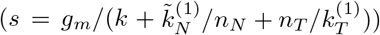. This is in support of a wide array of regulatory strategies observed in nature and will be discussed in more detail in the next section.

## 3 Discussion

Our first main result is that, contrary to intuition, toxin producers can promote diversity and act as the keystone species. This increase in diversity, however, comes at the cost of lower overall biomass and is a result of the trade-off between toxin resistance and nutrient competence [42, 26]. Second, we show that even when the toxins are not constitutively expressed but are regulated in response to the external environment, toxin producers can sustain diversity by creating a dynamic environment with persistence (oscillations and chaos) in species abundances.

### Shielding, tradeoffs, and competitive exclusion

To our knowledge, the shielding effect due to trade-offs involving toxins has not been previously explored. But similar mechanisms were suggested as a pathway to diversity for competitions over alternate nutrients [32]. In a nutshell, the shielding effect for alternate resources results from a shared enzyme budget, which flattens the fitness landscape where different strains grow at the same rate but use different nutrients [52, 43, 32, 53]. As for the case of toxins, the mechanistic basis for this tradeoff in some examples has been linked to the permeability of the cell membrane to both nutrients and toxins [42].

This idea goes beyond ecology. In adaptive immunity, trade-offs have been evoked to explain the diversity observed in *T* cells [2]. More generally, this is related to multi-objective optimization under a set of constraints, leading to a set of solutions that are equally optimal [51]. In the context of toxins, the trade-off is reflected in the balance between the ability to self-preserve against toxins and the ability to compete for nutrients [26, 42]. Interestingly, experiments show plasticity exhibited by these trade-offs in different environments [42, 25], *i*.*e*. the exact quantitative trade-off is environment dependent. Despite the seeming differences of the measured shapes, such that a convex (concave) trade-off would generally promote the fitness of the specialists (generalists) [12], our approach shows that both generalists and specialists share the same fitness under the shielding effect: for different values of *q* in Eq. (8) that determine these shapes, fitness equalization through the shielding effect remains valid for all species.

*Competitive exclusion* Generally, to reach steady-state coexistence in a flow driven system (for example, a chemostat) in Eq. (5), all surviving species reach the same growth rate that matches the flow rate 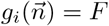 with *i* = 1,, *M*, where *M* is the total number of possible species in the system. Therefore, the solution to this system involves solving *M* equations with 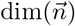 abiotic variables. This imposes an upper bound on the total number of species 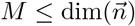 that may survive in the system and is referred to as the competitive exclusion principle [21]. Previous work exploring how toxins can increase diversity [48, 4] included the role of toxins by treating growth rates as functions of the various species abundances. Consequently, these systems are no longer limited by the competitive exclusion principle due to the increase in dimensionality that scales with the number of species, *i*.*e*. now the dimensions are 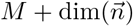. Here, rather than postulating a (ad hoc) growth function in the presence of toxins, we built a growth function using recent models for bacterial growth [57, 50], where growth depends only on available nutrients and the toxins produced but not on abundances of other species, making the theoretical more challenging but also more generally applicable.

Previous work has shown that cross-feeding through secreted metabolites can enhance diversity in conditions limited by nutrients [34, 37]. These secreted metabolites, unlike toxins, create commensal and mutualistic interactions. We here suggest that perhaps a community with a large variety of secreted toxins can play a similar role, even if its effect on diversity is not as intuitive as cross-feeding. The impact of toxin-mediated interactions between strains on diversity was explored using random networks [38]: although it was suggested that self-inhibition of toxin producers can contribute to diversity, toxin-mediated interactions in general decrease the likelihood for a community to diversify, as trade-offs were not considered in these models.

### Regulating toxins and persistence

Toxins are costly for producers and, therefore, are often regulated. However, the regulatory logic is not straightforward and shows a variety of strategies. For nutrient responses, although it is a common strategy that an increase in ppGpp, which is a signaling molecule in stringent responses such as nutrient limitation, up-regulates toxin production, there are also some cases where ppGpp suppresses toxin synthesis [7]. In response to toxins, autoinduction [49] senses toxins released through the lysis of sibling cells, which serves as an alarm cue of threats from its competitors. Hence, when the toxin level in the environment is elevated, an offensive approach is taken by producing more toxins. By responding to envelope stress, oxidative stress, and DNA damage through mechanisms such as the SOS response (which is important for DNA repairs) [7, 16], bacterial cells can, in effect, regulate toxin production through positive sensing. On the other hand, the same antibiotic stress can up-regulate or down-regulate the expressions of various type II toxin-antitoxin systems [40]. In our model of both nutrient sensing and toxin sensing, we see two different schemes that allow the toxin producer to survive: either large *σ* with positive sensing *p* > 0, or small *σ* with negative sensing *p* < 0. This seems to agree with the observed plethora of sensing schemes that act in opposite directions.

Our result shows that regulation of toxin production and its associated costs can change growth rates (fitness), allowing toxin producers to survive and potentially changing the overall diversity of the community. Here, however, the species can also be sustained in the system away from steady state, with species abundances showing persistence (oscillations and/or chaos). This is conceptually related to previously suggested resource competition models where oscillations are generated by competitions over at least three or more resources and is referred to as supersaturation [24, 23, 47, 14, 11]. Similarly, we show that persistence resulting from toxin regulation can also support a diversity higher than the number of abiotic factors and allow the system to tolerate the inexactness in trade-offs.

### The rise and evolutionary stability of toxin producers

Given a population of multiple species, how does the strategy of toxin production arise? Taking into account the cost of toxin production, we used the toxin production fraction *f*_1_ - the fraction of nutrients sacrificed for toxin production - to parameterize the ZNGI of the producer. With a large *f*_1_, the producer can dominate the population provided that its initial abundance is high. But this is not evolutionarily stable: any new producers introduced into the system through mutation would have a low initial abundance. With a small *f*_1_, the newly introduced producer can consistently invade the population. However, in this regime, the toxin producer has a higher nutrient efficiency (small 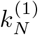) but lower toxin resistance (small 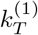) compared to the competing species that does not produce toxins. Does it even make sense for a toxin-sensitive species to produce that toxin into the environment, thereby lowering its competitiveness? Both large and small *f*_1_ align with the parameter space in [8] and [30], which consider only two-species competitions, and neither of the regimes reconciles with the rise of toxin production as a viable strategy. Building on their results, our work goes beyond two-species to multi-species competition, which provides a plausible explanation for this conundrum. In our model, it is still true that toxin production might not be a sensible strategy for specialists who invest in being the most nutrient-competent species. However, the toxin producer is now a keystone species, as we have shown in a high-diversity state: the community would collapse, and generalists that are neither the most nutrient-efficient nor the most toxin-resistant within the population would be driven to extinction if no toxin producers are present. By changing the production fraction such that *f*_*c*1_ < *f*_1_ < *f*_*c*2_ to shift the ZNGI, a generalist as a toxin producer can ensure its survival at a low abundance by sustaining a high diversity state through shielding. This is in support of the observed positive correlation between biodiversity and antimicrobial toxin genes abundances in rare taxa [33], and suggests that allelopathy might have a significant role in sustaining microbial diversity.

