## Supplementary Materials for "Promoting diversity in ecological systems through toxin production"

### Supporting Information Text

**Organization.** This document presents the derivation and justification of our choice of model.

**1. Derivation of growth rate functionals.** In this section of the SM, we show how we derived the growth rate functional and the production rate of toxins in a system with only one toxin and one resource.

***A. Growth rate with one nutrient and one toxin.*** Here is a quick recapitulation of the derivation of bacterial growth rate in (1, 2). At the exponential phase of bacterial growth, the growth rate $g$ is proportional to the amount of ribosomal proteins,

$$g = \frac{1}{M}\frac{dM}{dt} = \gamma\phi_R \; , \tag{S.1}$$

where $M$ is the total mass of protein and $\phi_R = M_R/M$ is the proteome fraction corresponds to ribosomal proteins with mass $M_R$. This is to account for the out-flux of the amino acid pool with size $I$ by translation in Fig. S1A. On the other hand, the in-flux of the pool is proportional to the amount of metabolic proteins with fraction $\phi_C$. At steady state, the size of the amino acid pool remains constant:

$$\frac{dI}{dt} = \nu\phi_C - \gamma\phi_R = 0 \; . \tag{S.2}$$

Here, $\nu$ and $\gamma$ denote the nutrient efficiency and translation efficiency respectively, which depend on the nutrients and toxins present in the environment and can be measured experimentally. In addition, the proteome constraint should be satisfied at any given time:

$$\phi_R + \phi_C = \phi \; , \tag{S.3}$$

where $1-\phi$ is the fixed fraction of the proteome that has no growth-rate dependency. This can be seen as a supply chain of proteins with two sub-processes: the upper stream process $C$ and the lower stream process $R$, carried out with the amount of resources $\phi_C$ and $\phi_R$ respectively. This is subjected to the constraint where the total amount of resources is fixed. By combining Eq. (S.2) with Eq. (S.3), we can solve for $\phi_R$, which in turn is related to growth rate by Eq. (S.1). This allows us to express growth rate in terms of the two efficiencies:

$$g = \frac{\phi}{\nu^{-1} + \gamma^{-1}} \; . \tag{S.4}$$

Suppose the external environment in Fig. S1A is nutrient-depleted, and that nutrient is brought in by transporter proteins, of which the amount scales with the proteome fraction $\phi_C$. This can be described by incorporating the Monod term in the nutrient efficiency $\nu$. Therefore, the inverse $\nu^{-1}$ is

$$\nu^{-1} = \nu_m^{-1}\frac{n_N + M_N}{n_N} \equiv \nu_m^{-1} + \frac{k_N^{(i)}}{n_N} \; . \tag{S.5}$$

The efficiency of the metabolic proteins is denoted by $\nu_m$, $M_N$ is the Monod constant, and $n_N$ is the nutrient level. We can further assume that translation is impaired by the toxin such that the translation efficiency decreases monotonically with the toxin level:

$$\gamma = \gamma_m \frac{M_T^q}{M_T^q + n_T^q} \; . \tag{S.6}$$

Therefore, the growth rate in Eq. (S.4) can be written as

$$g = \frac{g_m}{K + \frac{k_N}{n_N} + (\frac{n_T}{k_T})^q} \; , \tag{S.7}$$

with $K$ being a constant independent of the nutrient level and the toxin level.

***B. Production fraction in toxin production.*** We use the production fraction $f$ to take into account the cost of toxin production. It refers to the fraction of flux that is used to produce toxins. The steady-state equation in Eq. (S.2) is modified:

$$g = (1-f)\nu\phi_C = \gamma\phi_R \; . \tag{S.8}$$

Notice here that the rate of in-flux into $I$ in Fig. S1A is discounted by a fraction of $1-f$, or in other words, the growth rate can be rewritten as

$$g = \frac{g_m}{(1-f)^{-1}\nu^{-1} + \gamma^{-1}} \; . \tag{S.9}$$

Therefore, we modify the Monod constant as $k_N = \tilde{k}_N/(1-f)$. The flux is still conserved: the remaining fraction $f$ is diverted to producing toxins. The rate of toxin production per cell in Eq. (5) is modified as

$$\frac{\theta_T^+}{x} = f\nu\phi_C = f\nu\frac{g}{(1-f)\nu} = \frac{f}{1-f}g \; . \tag{S.10}$$

With constitutive production, the trade-off that allows shielding effect is

$$\frac{\tilde{k}_N}{1-f} + \frac{1}{k_T^q} = C \; . \tag{S.11}$$

Rearranging the terms gives

$$f = 1 - \frac{k_N}{C - 1/k_T^q} \; . \tag{S.12}$$

Under this trade-off, the production fraction $f$ increases (meaning that the species produces more toxins per cell) as $k_N$ decreases (species is efficient to grow on the resource) or $k_T$ increases (species is more resilient to the toxin). The gain in fitness in either nutrient efficiency or toxin resistance is balanced by the increasing cost of toxin production. This is graphically depicted in Fig. S2.

**2. Derivation for coexistence region.** In Sect. 2 of the main-text, we partition the abiotic factor space such that as the supply vector $(a_N, a_T)$ lies within region (ii) in Fig. 3, coexistence state is possible. Here we show the derivation. Dynamics of the toxin level is given by

$$\dot{n}_T = (a_T - n_T)F + \frac{f}{1-f} g_1 x_1 \; . \tag{S.13}$$

Setting $\dot{n}_T = 0$ at $g = F$, $x_1 = x_1^\star$ and $n_T = n_T^\star$ gives

$$\frac{f}{1-f} x_1^\star = n_T^\star - a_T \geq 0 \; . \tag{S.14}$$

The inequality comes from taking the abundance of the toxin producer to be non-negative at the coexistence solution, and this sets the upper bound of the toxin supply of region (ii): $a_T < n_T^\star$ .

How about the lower bound of the nutrient and toxin supply? We write down the dynamics of the nutrient level

$$\dot{n}_N = (a_N - n_N)F - g_1 x_1 - \sum_{i \neq 1} g_i x_i \; . \tag{S.15}$$

Substituting the expression of $x_1 = x_1^\star$ and $g = F$, the feasibility of the coexistence solution (which means the species abundances are non-negative) gives

$$\sum_{i \neq 1} x_i = (a_N - n_N^\star) - \frac{1-f}{f}(n_T^\star - a_T) \geq 0 \; . \tag{S.16}$$

Thus, we can recover the impact ratio $m_{T,N}^{(1)}$ of the producer species

$$\frac{n_T^\star - a_T}{n_N^\star - a_N} > \frac{1}{1 - 1/f} = m_1 \; . \tag{S.17}$$

If the coexistence is sustained by shielding effect, such that $g_i = F$, we can write down the condition as

$$\frac{1}{n_N} = n_T^q = \frac{g^m/F - K}{C} \equiv \frac{R}{C} \; , \tag{S.18}$$

where $C$ is defined in Eq. (S.11). In the main-text, we considered a change of variables with the appropriate scaling constants $L_N$ and $L_T$ for the nutrient and toxin levels:

$$\tilde{n}_T \to \frac{n_T}{L_T} \qquad ; \tilde{n}_N \to \frac{n_N}{L_N} \; . \tag{S.19}$$

with $R \equiv g_m/F - K$. The condition in Eq. (S.18) above becomes

$$\frac{1}{\tilde{n}_N L_N} = \tilde{n}_T^q L_T^q = \frac{R}{C} \; . \tag{S.20}$$

Therefore, we chose the length scales to be

$$\frac{1}{L_N} = L_T^q = \frac{R}{C} \; . \tag{S.21}$$

This gives $g(\tilde{n}_N = 1, \tilde{n}_T^q = 1) = F$. Therefore, for $q = 1$,the bound in Eq. (S.17) can be rewritten as

$$\frac{n_T^\star/L_T - a_T/L_T}{n_N^\star/L_N - a_N/L_N} > \left(\frac{R}{C}\right)^2 \frac{1}{1-1/f} = \left(\frac{R}{C}\right)^2 m_{T,N}^{(1)} \equiv m_1' \; . \tag{S.22}$$

which defines region (ii) and gives the lower bounds of the nutrient and toxin supplies. This gives us the impact line $n_T/L_T = m_i'(n_N/L_N - n_N^\star/L_N) + n_T^\star/L_T$.

**3. Physicality and stability of the coexistence solution.** Fig. 3 in Sect. 2 of the main-text shows two transitions: $f_1 = f_{c1}$ and $f_1 = f_{c2}$. What causes these transitions? The first transition is when the supply vector enters the region (ii) due to the increase of $|m^{(1)}_{T,N}|$, as proved in SM Sect. 2. There is a change of feasibility of the coexistence solution: the species abundances are non-negative when $f_1 > f_{c1}$. The second transition is caused by the change of local stability of the coexistence solution, as seen by the change of the largest eigenvalue of the Jacobian, from negative to positive. This means the coexistence solution is no longer stable at $f_1 > f_{c2}$. This is shown in Fig. S3

**4. Trade-off.** In the main-text, we evoke the trade-offs as a mathematical relation between $k_N^{(i)}$ and $k_T^{(i)}$, which describe nutrient competence and toxin resistance respectively. Can this relation be observed in the real system? In Fig. S4A, we adopt the data from (3) which shows the trade-offs between nutrient competence and resistance against three different antibiotics. In that study, Phan et al. look at the mechanistic structural root in cell permeability of engineered strains of *Escherichia coli* that gives rise to such antagonistic traits and attempt to describe the geometries of the trade-offs. They evaluate the toxin resistance of a strain by growing it on an agar plate with a gradient of the toxin level. They then evaluate the selection coefficient by comparing the growth of that strain in a chemostat under limiting nutrient conditions with that of a reference strain. It is suggested that there is a trade-off between the toxin resistance and the selection coefficient. As the permeability of the cell membrane increases, both nutrient uptake and antibiotic susceptibility increase due to passive diffusion. The consequence is that species that grow faster in a nutrient-limited environment would also have a lower toxin resistance; whereas species that grow slower would be more resilient in a toxic environment. It turns out that the shapes of trade-offs show great plasticity under different conditions (3, 4), and can be classified into three groups: decelerating, linear and accelerating. Thus, a valid theoretical model that explores trade-offs as a source of diversity should be able to accommodate and exhibit these different shapes and trends in the trade-offs.

The condition given by Eq. (6) in Sect. 2 of the main-text describes only the linear case. To describe the different shapes in the trade-offs, this can be generalized to

$$k_N^{(i)} + \left(\frac{1}{k_T^{(i)}}\right)^q = C\,, \qquad \forall i \tag{S.23}$$

where $C$ is a constant. Eq. (S.23) simply means that if species $i$ is well adapted to growing on the nutrient, with a small value of $k_N^{(i)}$, then it is more susceptible to the toxin with a small value of $k_T^{(i)}$. Using our model, we integrate the following system of equations:

$$\dot{x}_1 = \left(\frac{g_m}{1+(C-\left(1/k_T^{(1)}\right)^q)/n_N} - F\right)x_1\,; \tag{S.24}$$

$$\dot{x}_2 = \left(\frac{g_m}{1+k_N^{(2)}/n_N} - F\right)x_2\,; \tag{S.25}$$

$$\dot{n}_N = (a_N - n_N)F - \frac{g_m}{1+(C-\left(1/k_T^{(1)}\right)^q)/n_N}x_1 - \frac{g_m}{1+k_N^{(2)}/n_N}x_2\,. \tag{S.26}$$

up to some time $t = t_E$. This set of equations describes a system in the absence of toxins. The first line comes from writing down the dependency on nutrients as $k_N^{(1)} \equiv C - \left(1/k_T^{(1)}\right)^q$, which is the trade-off between toxin resistance with nutrient competence in Eq. (8). We can then consider some small perturbation $\epsilon$ from $(k_N^{(i)}, k_T^{(i)})$:

$$C = k_N^{(i)} + \frac{1}{(k_T^{(i)})^q} \tag{S.27}$$

$$= (1+\epsilon)k_N^{(i)} + \frac{1-\epsilon k_N^{(i)}(k_T^{(i)})^q}{(k_T^{(i)})^q}\,. \tag{S.28}$$

For example, as the membrane permeability decreases, the effective scale for the nutrient increases for species $j$ compared to species $i$ such that $k_N^{(j)} \equiv (1+\epsilon)k_N^{(i)}$. Then, by Eq. (S.28) the effective toxin level also increases. Effectively, the toxin resistance increases as the nutrient competence decreases. To quantitatively describe the nutrient competence of species $x_1$ in comparison with the reference, species $x_2$, we numerically integrate the system of equations with the initial conditions set to be $x_1(t=0) = x_2(t=0)$, and obtain the selection coefficient (5):

$$s \equiv \frac{\ln\frac{x_1(t_2)}{x_2(t_2)} - \ln\frac{x_1(t_1)}{x_2(t_1)}}{t_2 - t_1}\,. \tag{S.29}$$

This is a measurement that compares the exponential growth rates of a species $x_1$ and the reference species $x_2$. For comparison, we define the estimator of the selection coefficient to be

$$\hat{s} \equiv C_s\frac{\frac{g_m}{N^\star}(k_N^{(2)} - k_N^{(1)})}{(1+\frac{k_N^{(1)}}{N^\star})(1+\frac{k_N^{(2)}}{N^\star})}\,, \tag{S.30}$$

where $C_s$ is a scaling constant and

$$N^{\star} \equiv k_N^{\star(i)} \left(\frac{g_m}{F} - 1\right)^{-1} > 0 \; ; \qquad k_N^{\star(i)} \equiv \arg\min_i (k_N^{(1)}, k_N^{(2)}) \; . \qquad \text{[S.31]}$$

In other words, the estimator $\hat{s}$ is comparing the exponential growth rates in the absence of toxins, assuming the nutrient level is being dictated by the more nutrient-competent species. We then compare the selection coefficients $s$ and $\hat{s}$ of a species with how resistant it is against a toxin. We consider the following quantity as a proxy of toxin resistance:

$$s_t(k_T^{(1)}) \equiv 1 - 1/k_T^{(1)} \; . \qquad \text{[S.32]}$$

The interpretation of Eq. (S.32) is to take $IC_{50}$, which is the toxin level at which bacterial growth rate is halved, as an indicator of toxicity to the strain. The increase of $s_t$ reflects the increase of the effective scale for the toxin level $k_T^{(1)}$. Fig S4B-D shows that the prediction given by our estimator fits with the data from numerical integration, and exhibits three different trends as observed in the SPANC experiment in Fig. S4A.

The powerful implication of the constraint in Eq. (8) is that the shielding effect, which flattens the fitness landscape at the environmental condition in Eq. (S.33), is valid regardless of the shapes. At shielding, it does not particularly favor specialists (with extreme values of $k_N^{(i)}$ and $k_T^{(i)}$) as in the cases for convex trade-offs, nor generalists (with intermediate values of $k_N^{(i)}$ and $k_T^{(i)}$) as in the cases for concave trade-offs. Under this constraint, all species in the community can attain similar fitness at different values of $q$ if they can self-organize to change the environment such that

$$n_N = \frac{1}{n_T^q} \; . \qquad \text{[S.33]}$$

This is equivalent to the "shielding effect" in competition over alternative nutrient resources (or the term "shielded phase" in (6) which is also seen in (7)), which refers to self-organization of the microbes as a consortium to shields asymmetry of external supply of nutrients such that the re-scaled concentration is the same for all nutrients. We generalize this effect to toxins and trade-offs with different shapes (as captured by $q$).

**5. Effect of regulation on fitness.** What is the underlying mechanism behind sustaining diversity through positive sensing ($p > 0$) and negative sensing ($p < 0$)? The answer is that they both down-regulate toxin production (small production fraction $f_1$) within a wide range of toxin levels in the environment, thus lowering the production cost per cell. This can be seen from the phase diagram in Fig. 5, where the axis for the sensing scale $\sigma$ is in log-scale: there is a separation of scale in $\sigma$ for the coexistence region when $p > 0$ compared to that when $p < 0$.

We consider the case in Fig. 5C, where the toxin producer loses to the toxin non-producer when there is no regulation. The ZNGIs $g_i(n_N, n_T) = F$ plotted in Fig. S5 show how the cost of fitness is changed by nutrient sensing. With constitutive production ($p = 0$), the producer requires a higher nutrient level $n_N$ and a lower toxin level $n_T$ to survive compared to that of the non-producer. Fitness is not equalized since the ZNGIs do not intersect, and the producer is out-compete leading to a low diversity. However, for both regulation schemes ($p > 0$, $p < 0$), coexistence is possible either at large or small $\sigma$, as shown in Fig. 5C. This is because the producer is now fitter, and the required nutrient level for the producer to survive is lowered as shown in Fig. S5, which is a direct result from less nutrients being redirected from growth to toxin production. Fitness of the two species is equalized at the intersection of the ZNGIs, leading to either coexistence or persistence, depending on local stability. This sustain diversity that would otherwise collaps at constitutive regulation.

**6. Common mechanism for nutrient and toxin sensing .** For comparison with nutrient sensing in Fig. 5 in Sect. 2 of the main-text, we plotted the phase diagrams of toxin sensing ($s = n_T$ in Eq. (9)) for zone (2) and (3) in Fig. S6A-B. This shows that under both mechanisms, the behaviours of the systems are qualitatively the same. The similarities between nutrient sensing and toxin sensing suggest that both share a common mechanism; perhaps through sensing environmental cues, such as nutrient levels and toxin levels, the producer infers its competitiveness to decide whether to produce toxins.

To test this hypothesis, we set up a two-species system with Eq. (9) where the producer uses its growth rate when not producing toxins as a cue for regulation: $s = g_m/(k + \tilde{k}_N^{(1)}/n_N + n_T/k_T^{(1)})$. We call this growth rate sensing. This sensing is only a proxy for their competitiveness as $s$ does not depend on $f_1$: the sensing does not take into account the cost of toxin production. The phase diagrams of growth rate sensing for zone (2) is plotted in Fig. S6B and zone (3) in the in Fig. S6C-D. With the exception that it has an additional phase that shows bistability (T/C), growth rate sensing shows striking similarities compared to both nutrient sensing and toxin sensing, suggesting that inferring competitiveness is the unified mechanism behind both sensing schemes.

**7. Fine-tuning in supersaturation.**

***A. Supersaturation does not require fine-tuning of chemostat.*** In Sect. 2, we showed that regulation can sustain persistence which allows supersaturation. But is this a general behaviour, or does it require fine-tuning of the model parameters? In a chemostat model, there are three parameters that we can change externally: the flow rate ($F$), and the concentrations of nutrient and toxin ($a_N$,$a_T$) in the fresh medium pumped into the system. We considered adding a third species (green) to the system in Fig. 6 with the original two other species (blue, orange), and varied these parameters to obtain the phase diagrams in Fig. S7.

Within the chosen range of parameters, we see three phases: (P) Blue and orange species persist in limit cycle while green species is driven to extinction; (S) supersaturation of three species and; (I) Green species successfully invades and dominates. The considerable area occupied in the phase diagrams that correspond to phase (S) suggests that fine-tuning of the environment is not required for supersaturation. This agrees with supersaturation in system models with alternate resources (8).

***B. Supersaturation of four species.*** While supersaturation does not require the fine-tuning of parameters that define the environment, it does require fine-tuning in the phenotype of species. As shown in Sect. 2, supersaturation of three species can only be sustained within a margin of $\sim 1\%$ of the third species' toxin susceptibility $k_T^{(3)}$. In Fig. S8, we added to this system a fourth species and show the whole range of $k_T^{(3)}$ ($\sim 0.1\%$) that can sustain the survival of all four species. Supersaturation is not sustained outside of this range.

**8. Parameters.** Parameters for the figures that involve a single toxin producer are tabulated in Table. S1. For Fig. 5 and Fig. S6, we set $k_T^{(1)} = (1/(C - \tilde{k}_N^{(1)}/(1 - f_m/2)))$ (except Fig. 5C) and $k_T^{(2)} = (1/(C - \tilde{k}_N^{(2)}))$. For 5C, the constant $C$ is notated by ($\star$) because we set $k_T^{(1)} = (1/(C - \tilde{k}_N^{(1)}))$, meaning the trade-off is no longer valid at constitutive production. The length scales $L_N$ and $L_T$ are determined by Eq. (S.21) in SM. Sect. 2 in terms of $g_m$, $F$ and $C$. Next, we tabulated the parameters for the figures related to supersaturation through persistence in Table. S2. The scale of sensing is set as $\sigma = C/(g_m/F - K)$.

**Table S1. Parameters for the model with both constitutive and regulated production of toxins**

| Fig. | $g_m$ | $F$ | $\tilde{k}_N^{(1)}$ | $\tilde{k}_N^{(2)}$ | $\tilde{k}_N^{(3)}$ | $\tilde{k}_N^{(4)}$ | $\tilde{k}_N^{(5)}$ | $C$ | $K$ | $q$ | $f_m$ | $a_N$ | $a_T$ |
|---|---|---|---|---|---|---|---|---|---|---|---|---|---|
| 3 | 1.2 | 1.02 | 0.2 | 0.35 | 0.5 | 0.65 | 0.8 | 3 | 0.1 | 1 | - | 10 | 0 |
| 4B | 1.2 | 1.03 | 0.1 | 0.6 | 0.7 | 0.8 | 0.9 | 5 | 0 | 1 | 0.1 | 14 | 0 |
| 5A-B | 1.2 | 1.03 | 0.4 | 0.6 | - | - | - | 2 | 0.1 | 1 | - | 14 | 0.1 |
| 5C, S5 | 1.2 | 1.03 | 0.4 | 0.6 | - | - | - | 2 ($\star$) | 0.1 | 1 | 0.9 | 14 | 0.1 |
| 5D | 1.2 | 1.03 | 0.6 | 0.4 | - | - | - | 2 | 0.1 | 1 | 0.9 | 14 | 0.1 |
| S6 | 1.2 | 1.03 | 0.4 | 0.6 | - | - | - | 2 | 0.1 | 1 | - | 14 | 0.1 |

**Table S2. Parameters for our model showing supersaturation due to persistence**

| Fig. | $g_m$ | $F$ | $\tilde{k}_N^{(1)}$ | $\tilde{k}_N^{(2)}$ | $\tilde{k}_N^{(3)}$ | $\tilde{k}_N^{(4)}$ | $\tilde{k}_T^{(4)}$ | $C$ | $K$ | $q$ | $f_m$ | $\sigma$ | $p$ | $a_N$ | $a_T$ |
|---|---|---|---|---|---|---|---|---|---|---|---|---|---|---|---|
| 6 | 1.2 | 1.03 | 0.4 | 0.6 | 0.41 | - | - | 2 | 0.1 | 1 | 0.55 | 1.87785 | 1.8 | 14 | 0.1 |
| S7A | 1.2 | 1.03 | 0.4 | 0.6 | 0.41 | 0.7 | 0.864583 | 2 | 0.1 | 1 | 0.55 | 1.87785 | 1.8 | - | - |
| S7B | 1.2 | $1.03 + \Delta F$ | 0.4 | 0.6 | 0.41 | 0.7 | 0.864583 | 2 | 0.1 | 1 | 0.55 | 1.87785 | 1.8 | 14 | 0.1 |
| S8 | 1.2 | 1.03 | 0.4 | 0.6 | 0.41 | 0.7 | 0.75 | 2 | 0.1 | 1 | 0.55 | 1.87785 | 1.8 | 14 | 0.1 |

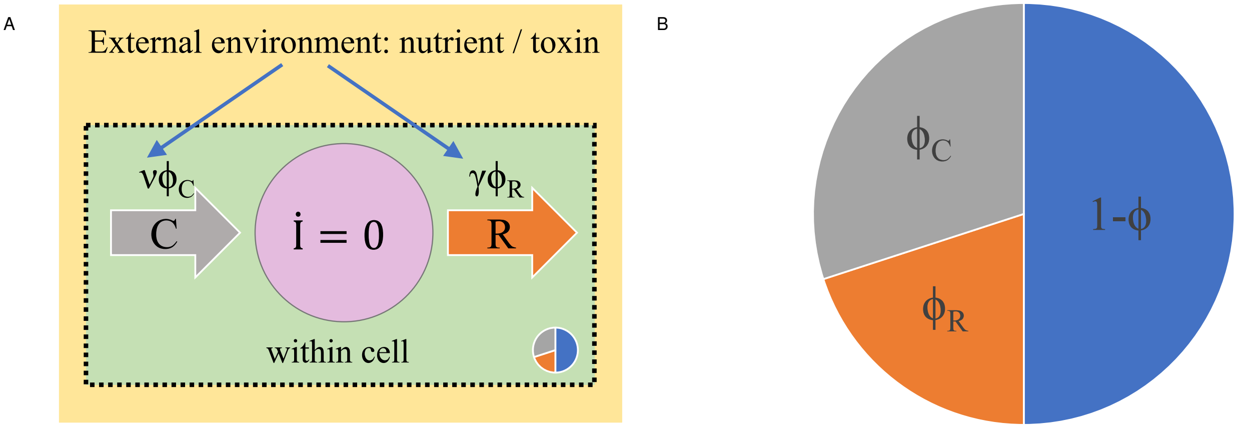


**Fig. S1.** Rate of bacterial growth depends on both proteome partitioning within the cell and abiotic factors in the external environment. (A) Suppose the rate of bacterial growth is limited by two sub-processes $C$ and $R$, contributing to the in-flux and out-flux of some intermediate, such as amino acids, with abundance $I$. The rate of the sub-processes is $\nu\phi_C$ and $\gamma\phi_R$. The rates depend on the efficiencies $\nu$ and $\gamma$ which are affected by nutrient and toxin levels in the external environment, and the fraction of corresponding proteome partition. (B) The fraction of proteome $\phi = \phi_C + \phi_R$ associated with growth is fixed and is referred to as the proteome constraint. The sub-fractions $\phi_C$ and $\phi_R$ associated with sub-processes $C$ and $R$ vary to ensure the steady state of the intermediate $\dot{I} = 0$, as seen in (2).

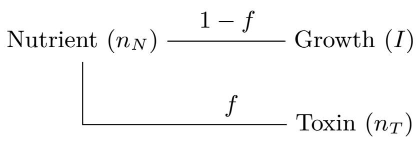


**Fig. S2.** Flux conservation in toxin production. We define production fraction $f$ which is the contribution of flux into toxin production. The remaining fraction $1 - f$ is used for bacterial growth, corresponding to $\dot{I} > 0$ in Fig. S1A. Toxin is produced at a cost of reduction of growth rate. Effectively, the production fraction modifies the Michaelis-Menton constant.

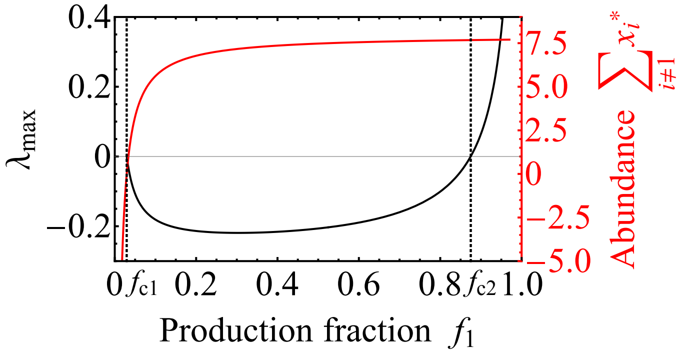


**Fig. S3.** The effect of increasing toxin production. The largest eigenvalue $\lambda_{max}$ (black) of the coexistence solution and the corresponding total abundance of all species excluding the producer (red) as production fraction $f_1$ changes, which is stable only when $f_{c1} < f_1 < f_{c2}$.

 

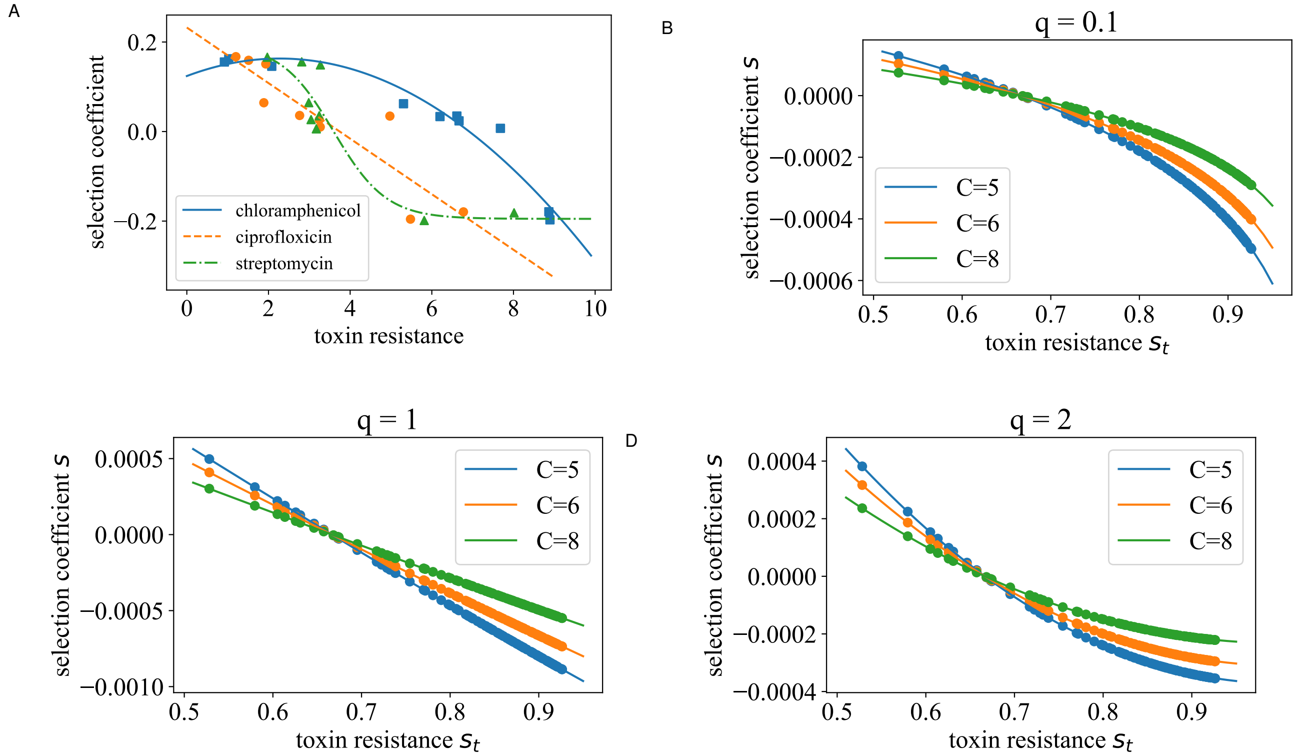


**Fig. S4.** (A) Data adapted from Ferenci et al (3). showing three different trends of nutrient competence as toxin resistance increases. These trends are qualitatively captured by the model: (B) accelerating (C) linear (D) decelerating. Solid lines are theoretical differences in fitness $\hat{s}$ with $C_s = 0.1$, and data points are selection coefficient $s$ obtained by numerical integration with parameters drawn from sampling.

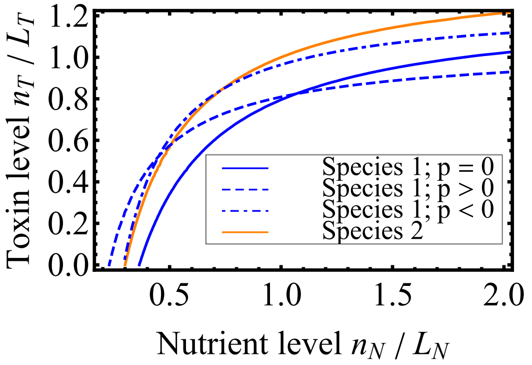


**Fig. S5.** Comparison of constitutive production with regulation production through nutrient sensing with one producer corresponding to Fig. 5C. ZNGIs for the toxin producer (blue) and non-producing species (orange).Dash line refers to positive sensing ($p = 1.2$, $\sigma = 2$) and dash-dotted refers to negative sensing ($p = -1$, $\sigma = 0.3$). Both lead to the coexistence of the toxin producer and non-producer.

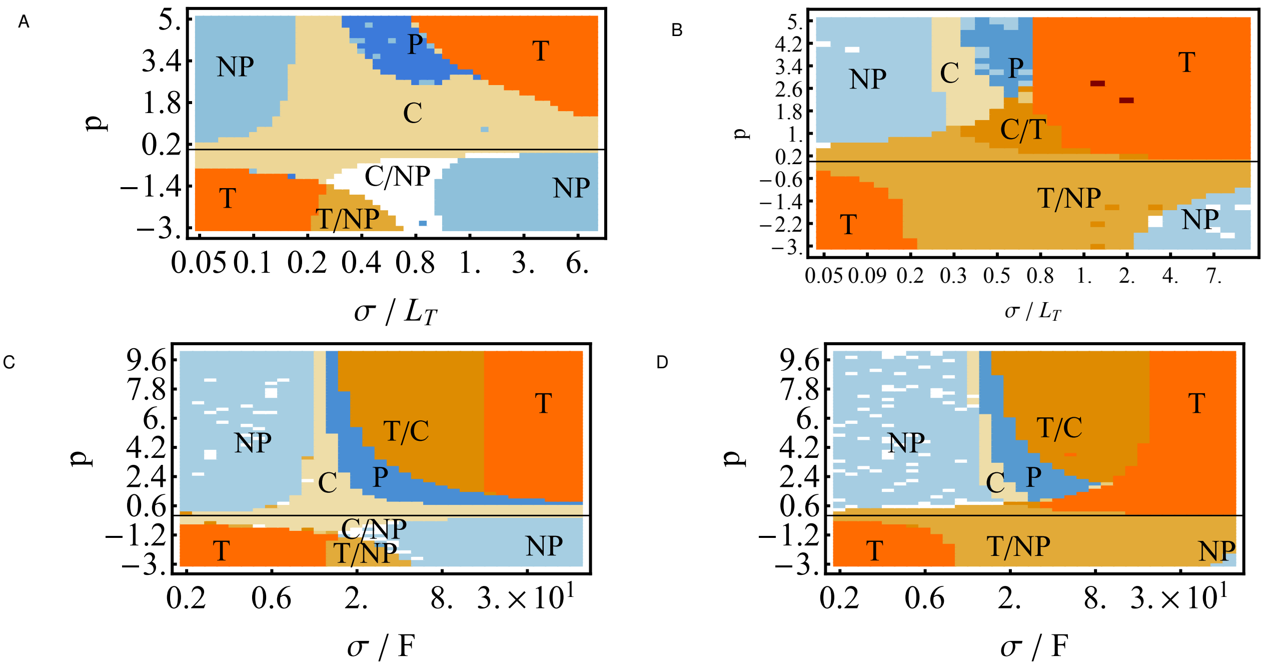


**Fig. S6.** Phase diagrams with varying Hill function $p$ and scaling constant $\sigma$ at zone (2) with regulation through (A-B) toxin sensing and (C-D) growth rate sensing, with constant $L_T$ and flow rate $F$ setting the scales. The maximum ratios $f_m$ are 0.55 (A,C) and 0.9 (B,D).

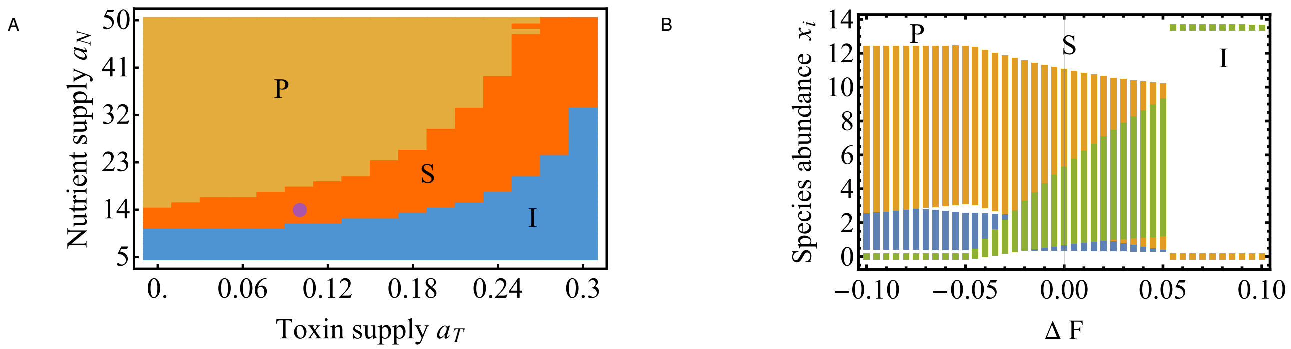


**Fig. S7.** Supersaturation of three species with one nutrient and one toxin at varying (A) toxin and nutrient supply concentrations and (B) flow rate $F = 1.03 + \Delta F$, showing three different phases. P denotes the phase that the invader (green) fails to invade the persistence generated by the original pair of toxin producer (blue) and non-producer (green). I denote the population being dominated by the invader. S is supersaturation. The supply concentrations of (B) are denoted by the purple dot in (A).

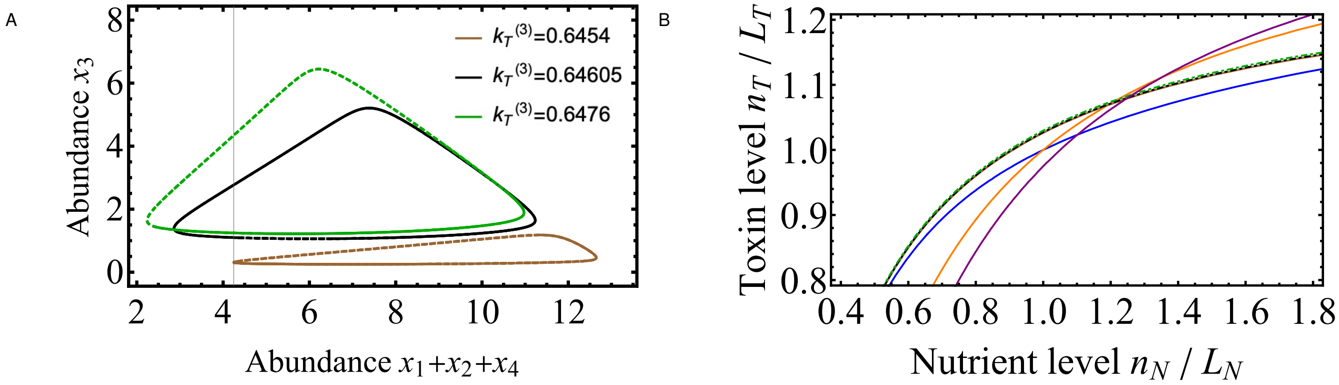


**Fig. S8.** Supersaturation of four species supported by a toxin and a nutrient source due to negative feedback under nutrient sensing. We vary the toxin resistance of the third species $k_T^{(3)}$ (Green, black and brown). (A) Limit cycles of species abundances after transient dynamics. Supersaturation is not sustainable outside of this range. (B) ZNGIs of the four species. Slight changes in toxin resistance of the third species greatly affect the fitness and the competition outcome.
